## Supplementary figures for "Ancient diversity in host-parasite interaction genes in a model parasitic nematode"

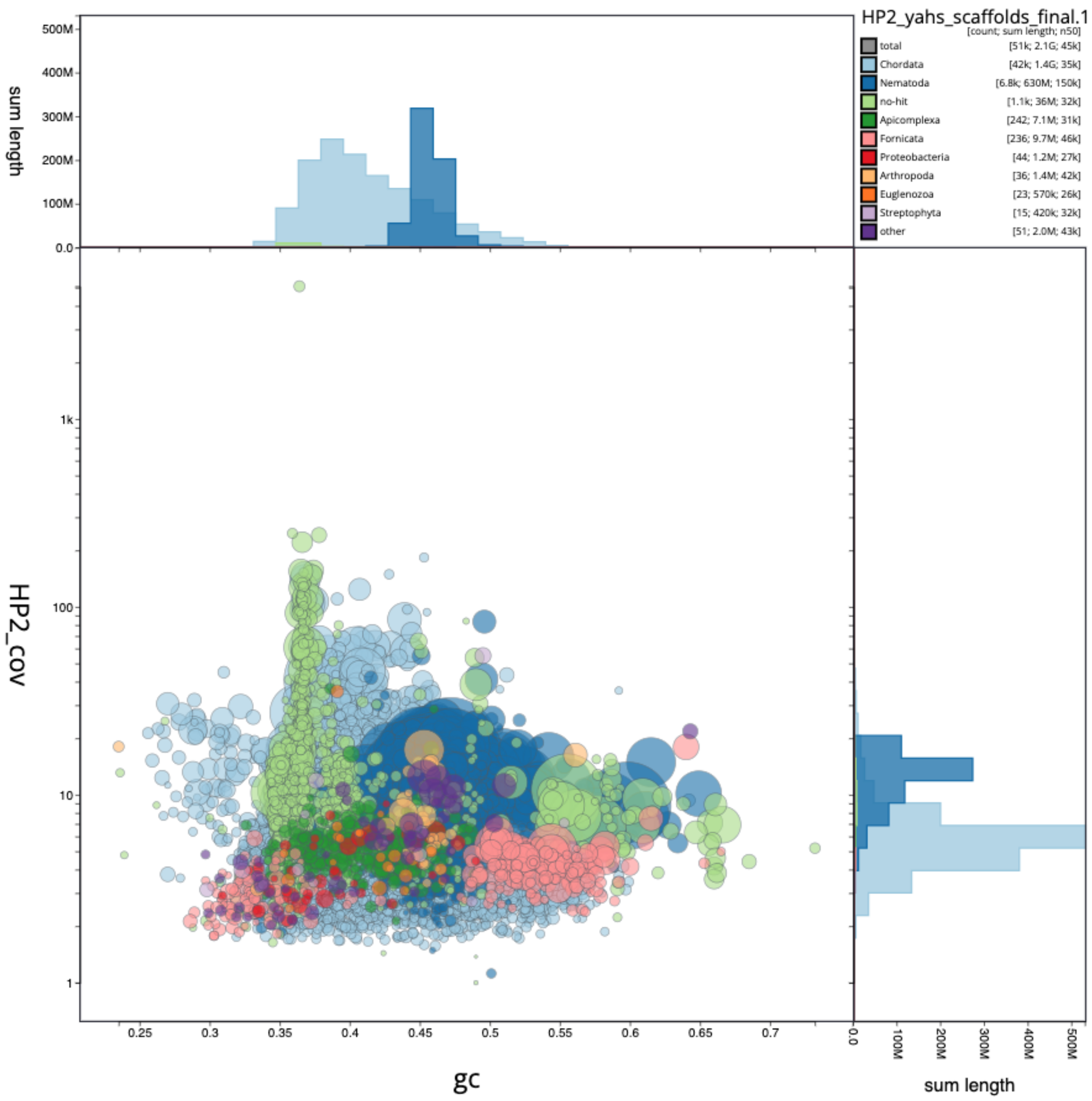


**Figure S1: Blob plot of *Heligmosomoides polygyrus* ngHelPoly2 dataset**

Blob plot of the assembly of the *H. polygyrus* ngHelPoly2 dataset showing extensive contamination by host and two diplomonad parasites (a *Giardia* sp. and a *Spironucleus* sp.). The x-axis represents GC content and the y-axis represents PacBio HiFi read coverage. Scaffolds are coloured by phylum. Circles are sized in proportion to scaffold length on a square-root scale, ranging from 11.3 to 86.5 kb. Histograms show the distribution of scaffold length sum along each axis. Two groups of scaffolds are likely derived from Fornicata (Metamonada) parasites: the leftmost blob corresponds to a *Spironucleus* sp. and the rightmost blob corresponds to a *Giardia* sp. Note that scaffolds labelled as apicomplexan are mislabelled *Apodemus sylvaticus* host scaffolds; all hits are to a *Plasmodium yoelii yoelii* genome in NCBI (GCA_000003085.2) which is contaminated with host rodent sequence.

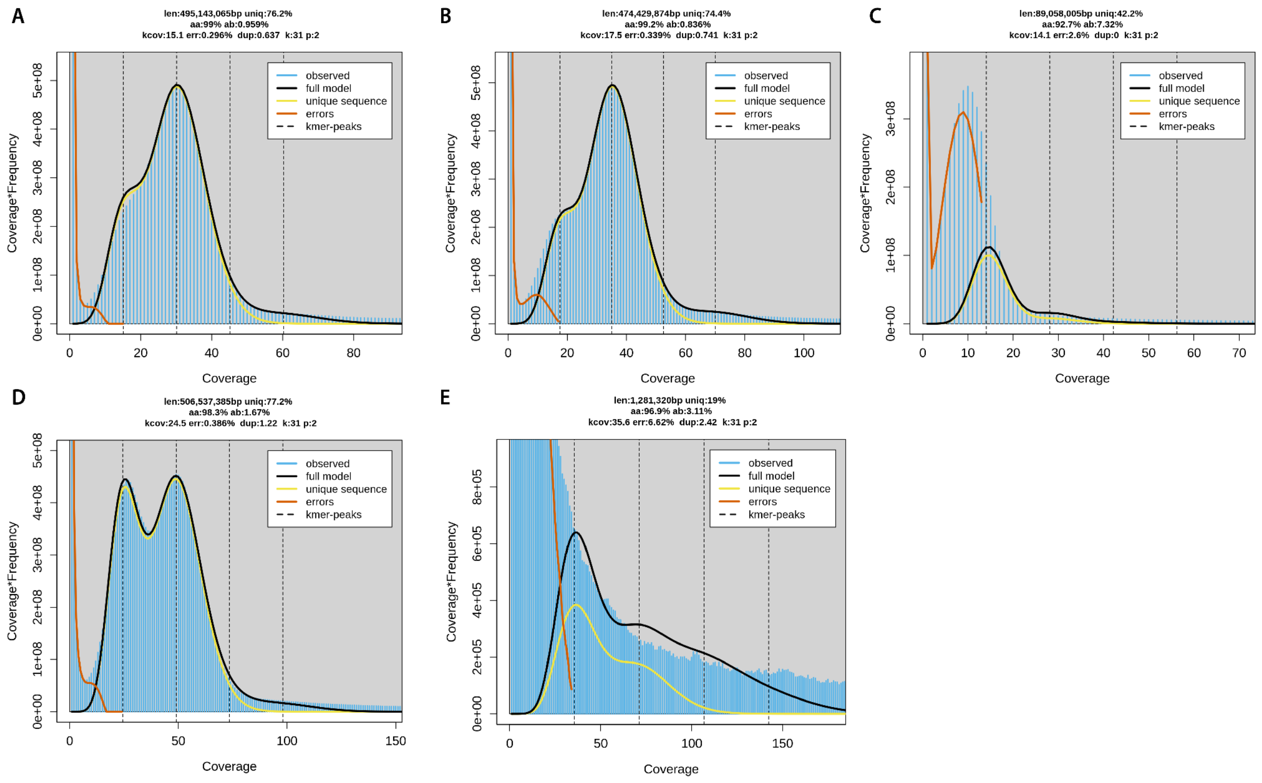


### Figure S2: *K*-mer profiles of PacBio HiFi data for all *Heligmosomoides bakeri* and *Heligmosomoides polygyrus* individuals

*K*-mer profiles (k = 31) and fitted models for PacBio raw HiFi read data from GenomeScope 2.0 for (A) *H. bakeri* nxHelBake1, (B) *H. bakeri* nxHelBake2, (C) *H. bakeri* nxHelBake3, (D) *H. polygyrus* ngHelPoly1 and (E) *H. polygyrus* ngHelPoly2. The *k*-mer profile shown for *H. polygyrus* ngHelPoly2 is of the read set after removing reads that map to *Apodemus sylvaticus* and *Giardia muris*. The genome size and heterozygosity estimates for *H. bakeri* nxHelBake3 (C) and *H. polygyrus* ngHelPoly2 (E) are not reliable due to low coverage. In the high coverage datasets, we note that genome size estimates are substantially less than the assembled reference genomes, which may be caused by high copy number *k*-mers being poorly modelled.

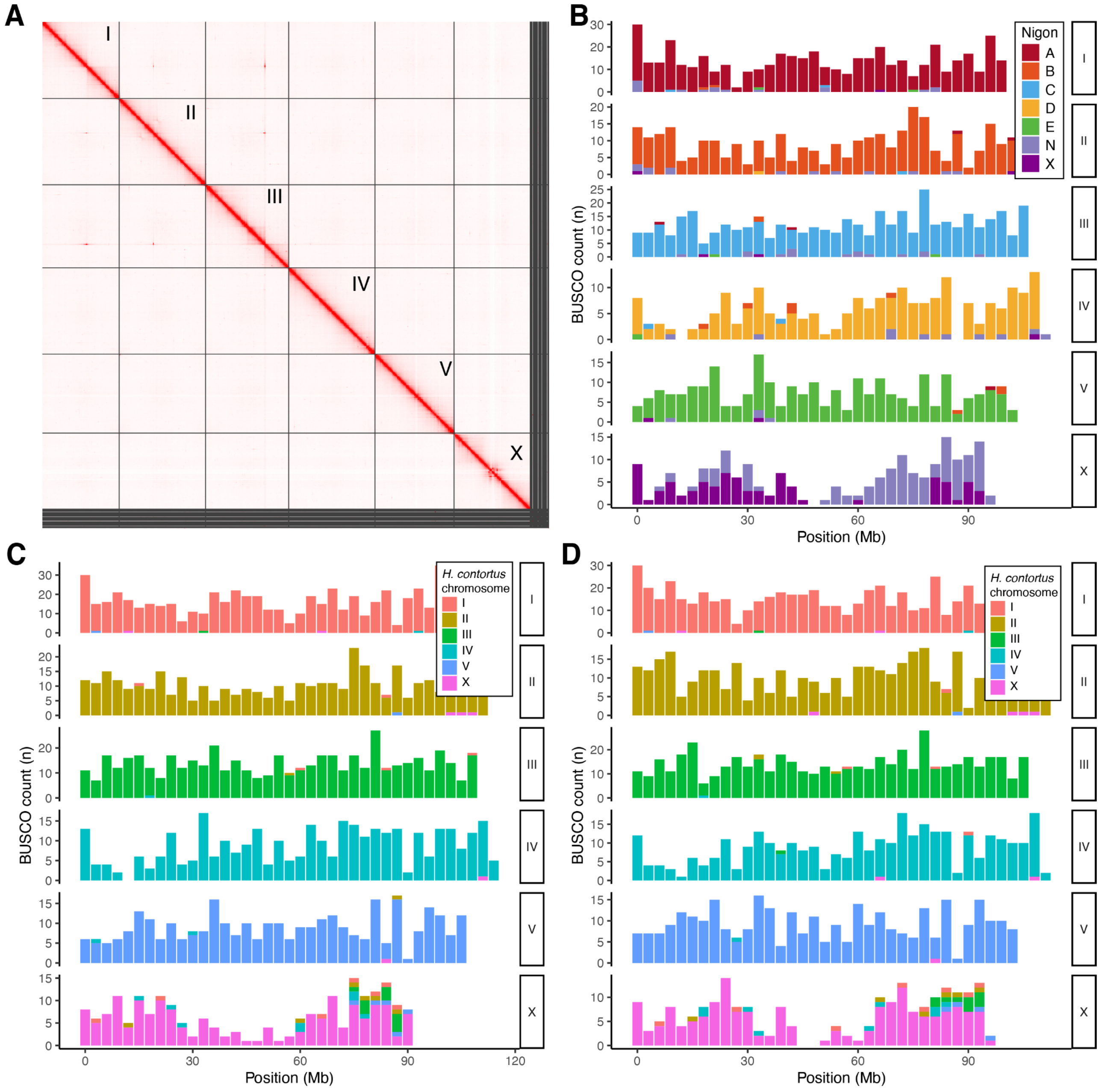


**Figure S3: Chromosome-level reference genomes for *Heligmosomoides bakeri* and *Heligmosomoides polygyrus***

(A) Hi-C contact map for ngHelPoly1.1 reference genome. Chromosome names are indicated. (B) Distribution of BUSCO genes in the six ngHelPoly1.1 chromosomes coloured by Nigon element (Gonzalez de la Rosa et al. 2021). Distribution of BUSCO genes coloured by their location in the chromosome-level *H. contortus* (PRJEB506) genome in the (C) *H. bakeri* nxHelBake1.1 and (D) *H. polygyrus* ngHelPoly1.1 reference genomes.

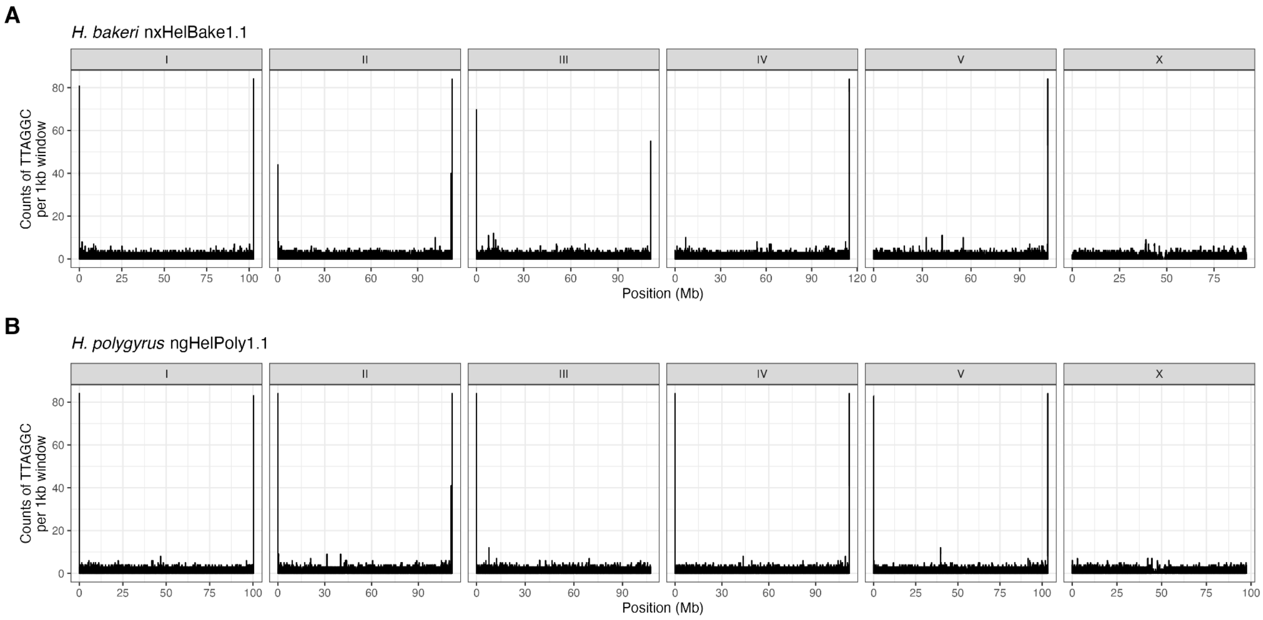


**Figure S4: Telomeric repeat sequence in *Heligmosomoides* reference genomes**
Counts of the nematode telomeric repeat sequence (TTAGGC) in 1 kb windows in the (A) *H. bakeri* nxHelBake1 and (B) *H. polygyrus* ngHelPoly1 reference genomes. Telomeric repeat counts are shown for the six chromosome-sized scaffolds only.

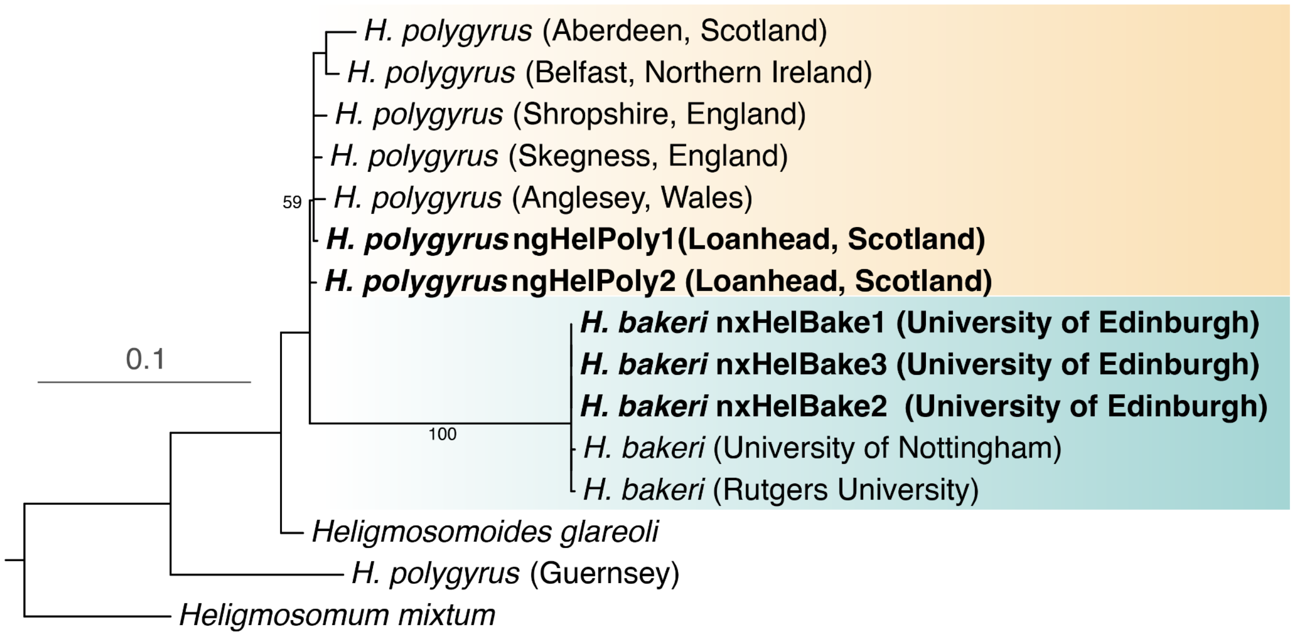


**Figure S5: Cytochrome oxidase 1 (COI) phylogeny of Heligmosomoides and related nematodes**

Maximum likelihood phylogeny of the mitochondrial cytochrome oxidase 1 gene in laboratory isolates of *H. bakeri*, wild isolates of *H. polygyrus*, and outgroup taxa. The sequences are derived from (Cable et al. 2006) or the mitochondrial genomes of individuals sequenced as part of this work (highlighted in bold). The origin of each isolate is shown in parentheses. Bootstrap support values are shown for the branches subtending the *H. bakeri* and *H. polygyrus* clades. Branch lengths represent the number of substitutions per site; scale is shown. As noted by (Cable et al. 2006) and (Maizels et al. 2011), the COI sequence from the "*H. polygyrus*" isolate from Guernsey is highly divergent from other *H. polygyrus* isolates. We believe this is caused either by misidentification or a low-quality sequence.

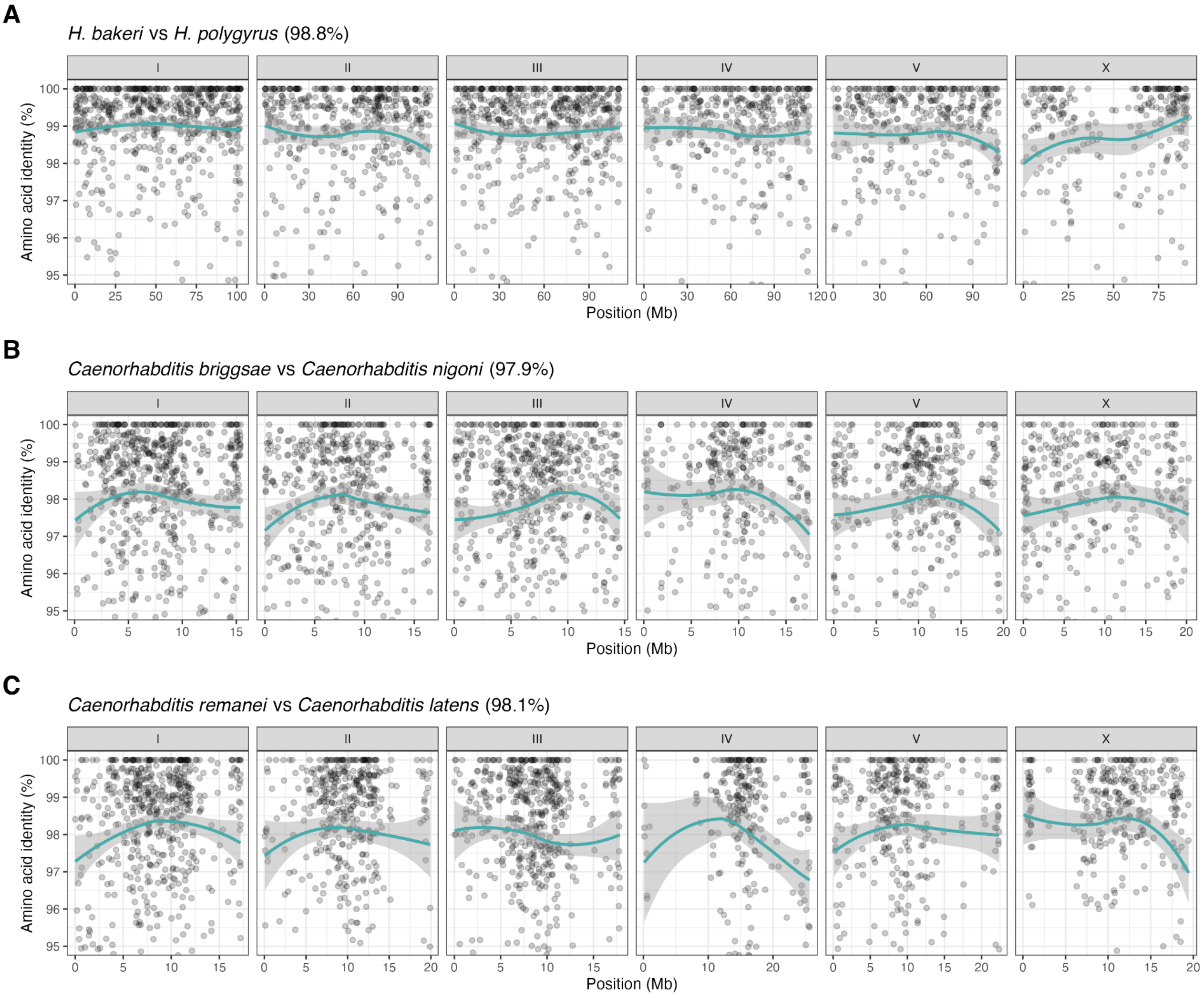


**Figure S6: Identity of highly conserved proteins in three nematode species pairs**
Protein identity of BUSCO proteins in (A) *H. bakeri* and *H. polygyrus* (2,667 proteins) (B) *Caenorhabditis briggsae* and *Caenorhabditis nigoni* (2,923 proteins) and (C) *Caenorhabditis remanei* and *Caenorhabditis latens* (2,532 proteins). Lines represent LOESS smoothing functions fitted to the data; standard error is shown using grey shading. The mean protein identity is shown in parentheses. BUSCOs were identified in isoform-filtered proteomes of each species using the protein mode and the nematoda_odb10 lineage. Protein sequences were aligned using FSA and proteins that had less than 85% identity were filtered out to avoid alignment errors biassing divergence estimates.

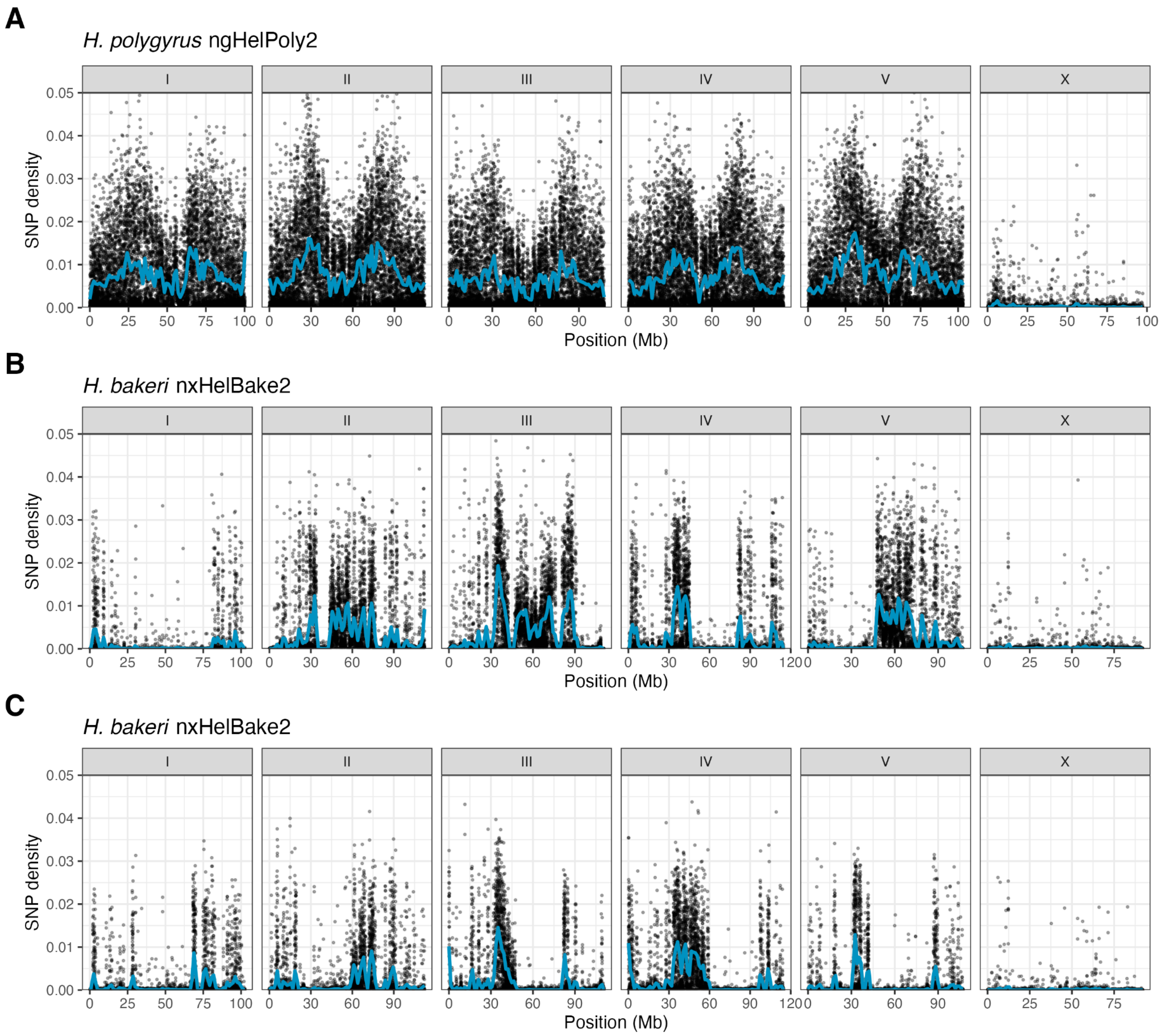


**Figure S7: Distribution of heterozygous SNPs in *Heligmosomoides bakeri* and *Heligmosomoides polygyrus***

Distribution of heterozygous SNPs in (A) *H. polygyrus* ngHelPoly2 relative to the ngHelPoly1.1 reference genome and the (B) *H. bakeri* nxHelBake2 and (C) *H. bakeri* nxHelBake3 relative to the nxHelBake1 reference genome. Points represent the density of biallelic SNPs in 10 kb windows. All three individuals are male and therefore the X chromosome is hemizygous. SNPs called in repeat-containing regions were filtered and calculated SNP density is calculated as the number of non-repetitive SNPs per non-repetitive base. Homozygous alternate variants (i.e. variants that represented differences from the reference genome rather than heterozygous SNPs) were ignored. Lines represent LOESS smoothing curves fitted to the data.

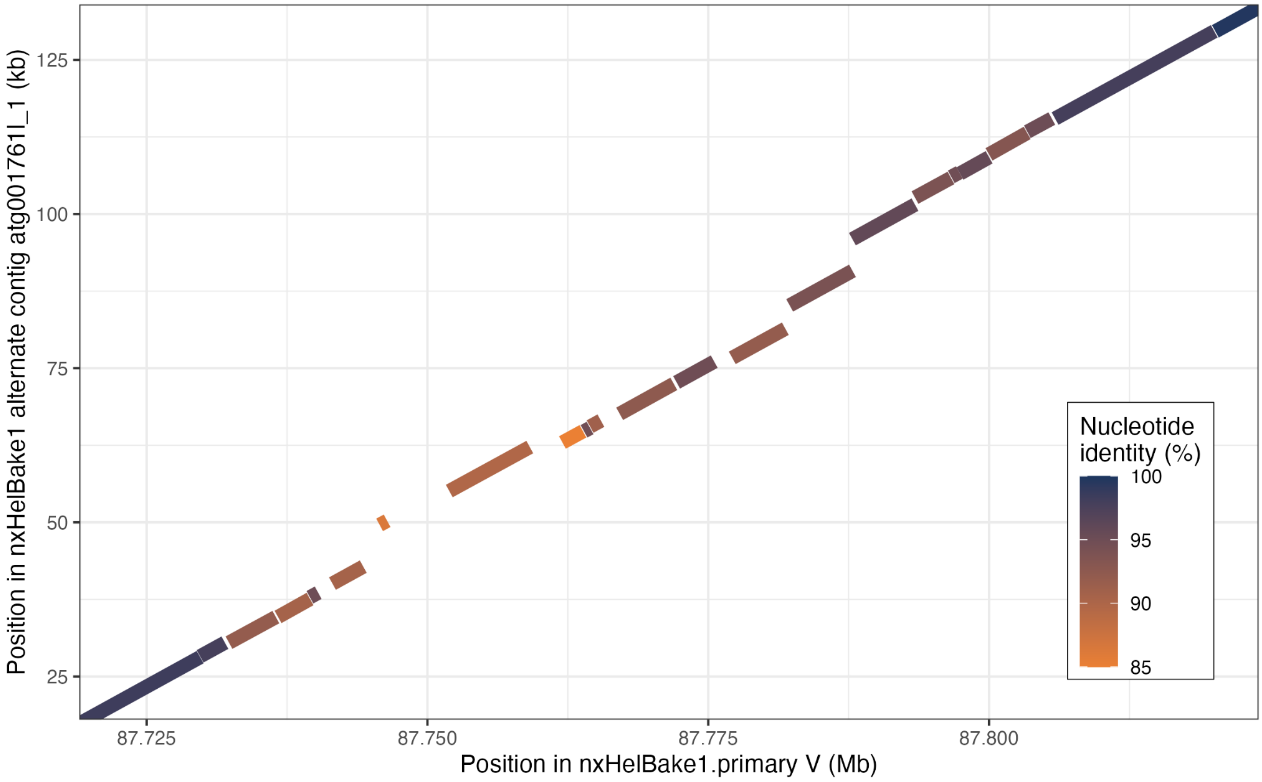


**Figure S8: Example hyper-divergent haplotype on *H. bakeri* nxHelBake1 chromosome V**

Nucleotide alignment between nxHelBake1 alternate contig atg001761l_1 and nxHelBake1 primary chromosome V (87.71 - 87.86) with each aligned segment coloured by its nucleotide identity. Repetitive alignments are not shown. The two non-divergent flanking alignments show high nucleotide identity (99.48% and 99.73%, respectively) whereas several aligned segments within the hyper-divergent haplotype have nucleotide identities of < 90%. Read alignments for this region are shown in Figure 3C.

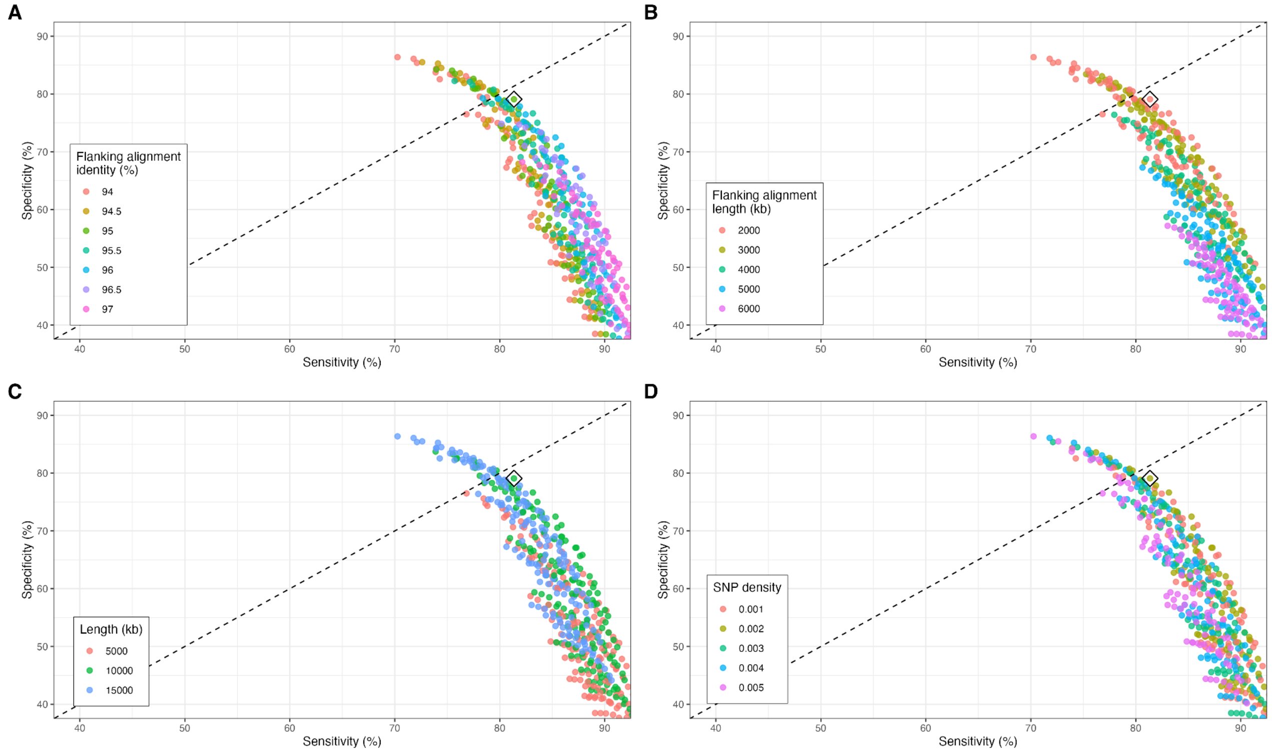


**Figure S9: Optimising the hyper-divergent region calling pipeline**

Each point represents an independent run of the hyper-divergent haplotype calling pipeline on PacBio CLR assemblies for 15 *C. elegans* strains (Lee et al. 2021) using a different parameter set. The output of each run is summarised by sensitivity on the X-axis (the number of overlapping bases divided by the total number of bases defined by (Lee et al. 2021)) and specificity (the number of overlapping bases divided by the total number of bases classified as hyper-divergent by our approach). Each panel represents different parameter: (A) flanking alignment identities of 94-97%, (B) flanking alignment lengths of 2-6 kb, C) minimum size of alignment gaps to be considered as a hyper-divergent haplotype of 5-15 kb, and (D) SNP density, derived from assembly-based variant calling, within alignment gaps of 0.001-0.005. The diamond represents the chosen parameter set.

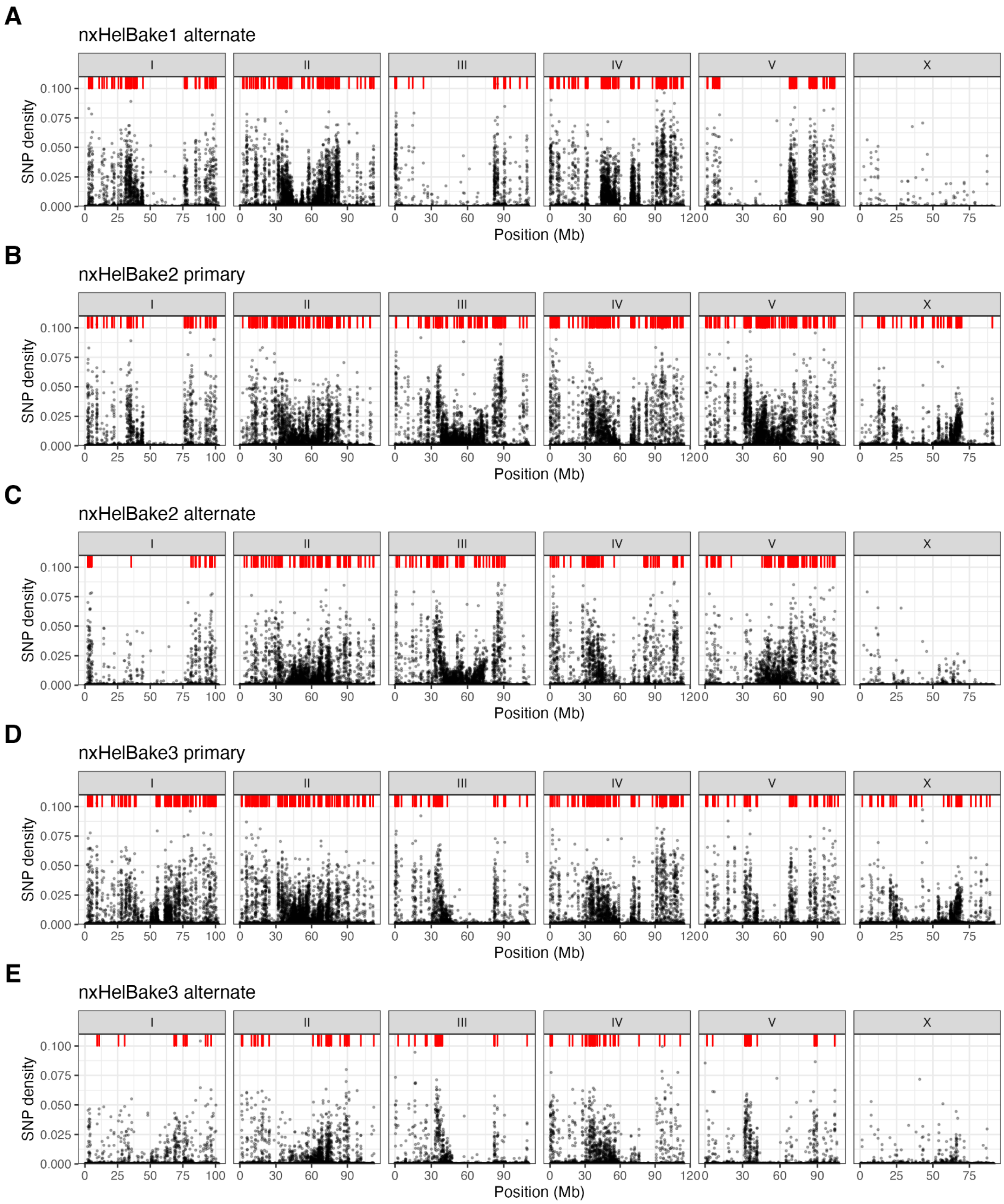


**Figure S10: Locations of hyper-divergent haplotypes across all three individuals**

The locations of hyper-divergent haplotypes in all five non-reference haplotypes: (a) nxHelBake1 alternate, (b) nxHelBake2 primary, (c) nxHelBake2 alternate, (d) nxHelBake3 primary, (e) nxHelBake3 alternate. Red boxes represent locations of hyper-divergent haplotypes. Hyper-divergent haplotypes called on the X chromosome in the three alternate haplotypes were removed (2, 1, and 1 from nxHelBake1, nxHelBake2, and nxHelBake3 alternate assemblies, respectively). The distribution of heterozygous SNPs, derived from assembly-based variant calling, in 10 kb windows are shown for each haplotype. SNPs that overlapped with a repeat annotation were removed and SNP density was calculated using the remaining SNPs and the number of non-repetitive bases in each window.

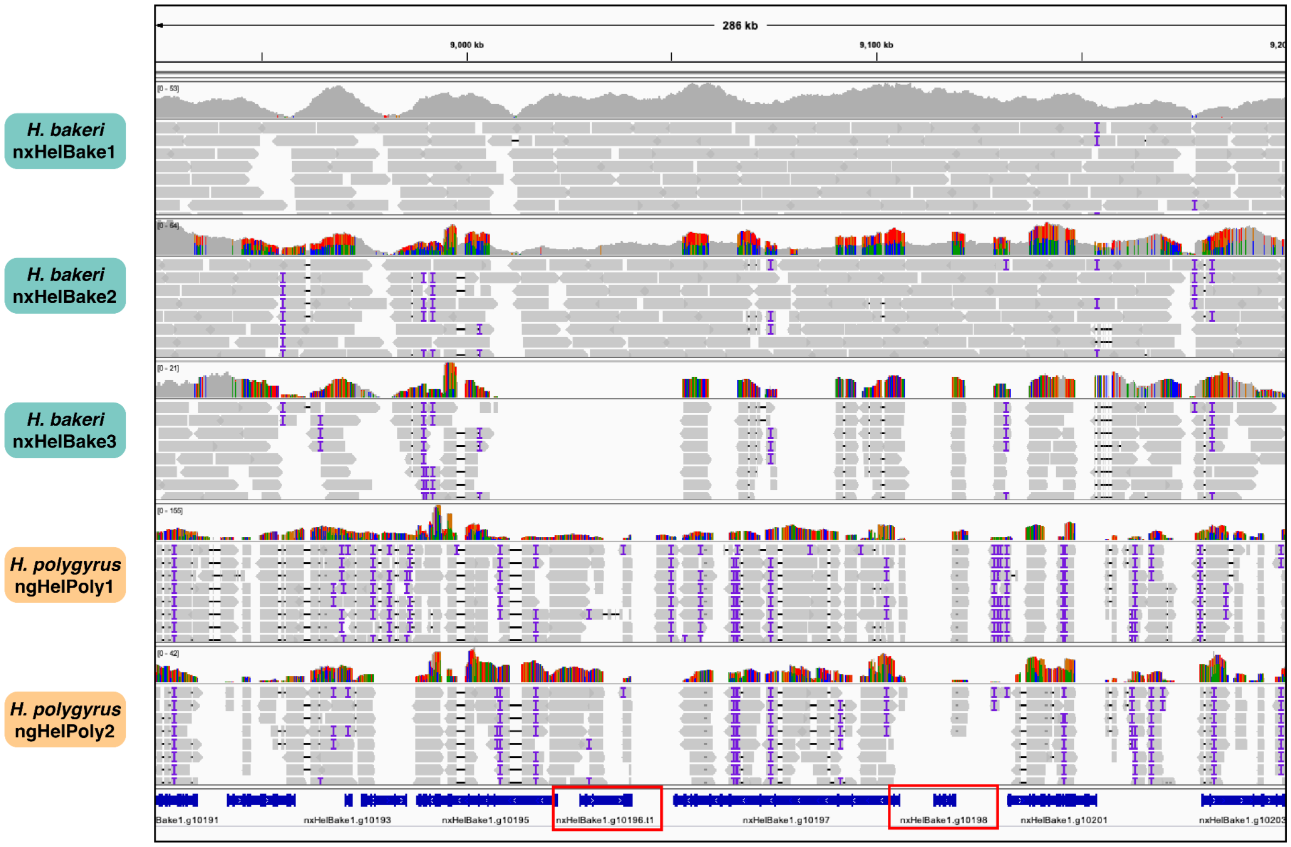


**Figure S11: Read alignments in a region in nxHelBake1.1 containing two *Ancylostoma*-secreted proteins showing evidence of trans-specific polymorphism**

PacBio HiFi read alignments of all sequenced individuals to the nxHelBake1.1 reference genome in a 287 kb region on chromosome I (I:8.92-9.19 Mb) showing evidence of trans-specific polymorphisms. Two genes in this region belong to the ASP family (nxHelBake1.g10196 and nxHelBake1.g10198; highlighted in red boxes). The top panel shows the coverage and the bottom panel shows aligned PacBio HiFi reads. The coloured vertical lines indicate mismatched bases at that position. Note that mismatched bases are only shown in the coverage tracks and not in the read alignments at this zoom level in IGV.

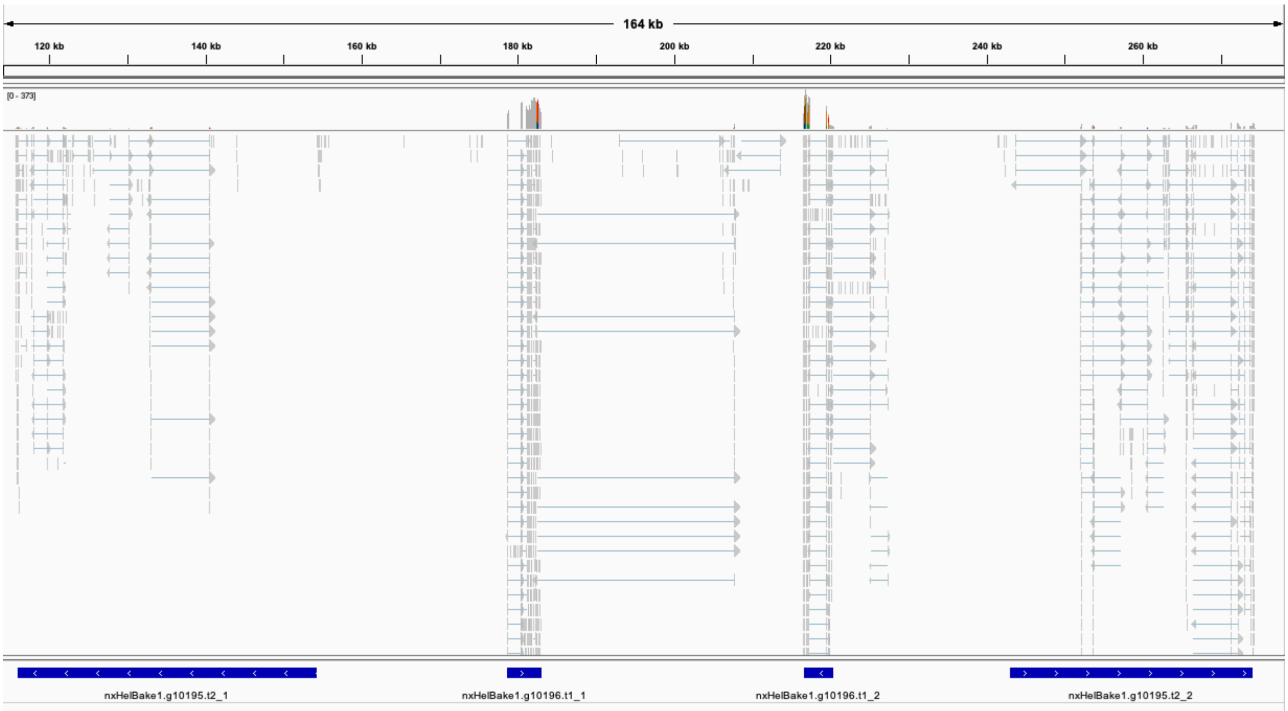


**Figure S12: RNA-seq support for genes predicted in the alternate *H. bakeri* haplotype**
Alignments of short-read RNA-seq data collected from pools of *H. bakeri* individuals (Rausch et al. 2018) to the alternate haplotype from the nxHelBake2 primary assembly. The top panel shows the coverage and the bottom panel shows aligned RNA-seq reads. Both homologues of nxHelBake1.g10195 (which contains a growth factor receptor domain) and both homologues of the ASP nxHelBake1.g10196 (a member of the *Ancylostoma*-secreted protein) are supported by RNA-seq reads.

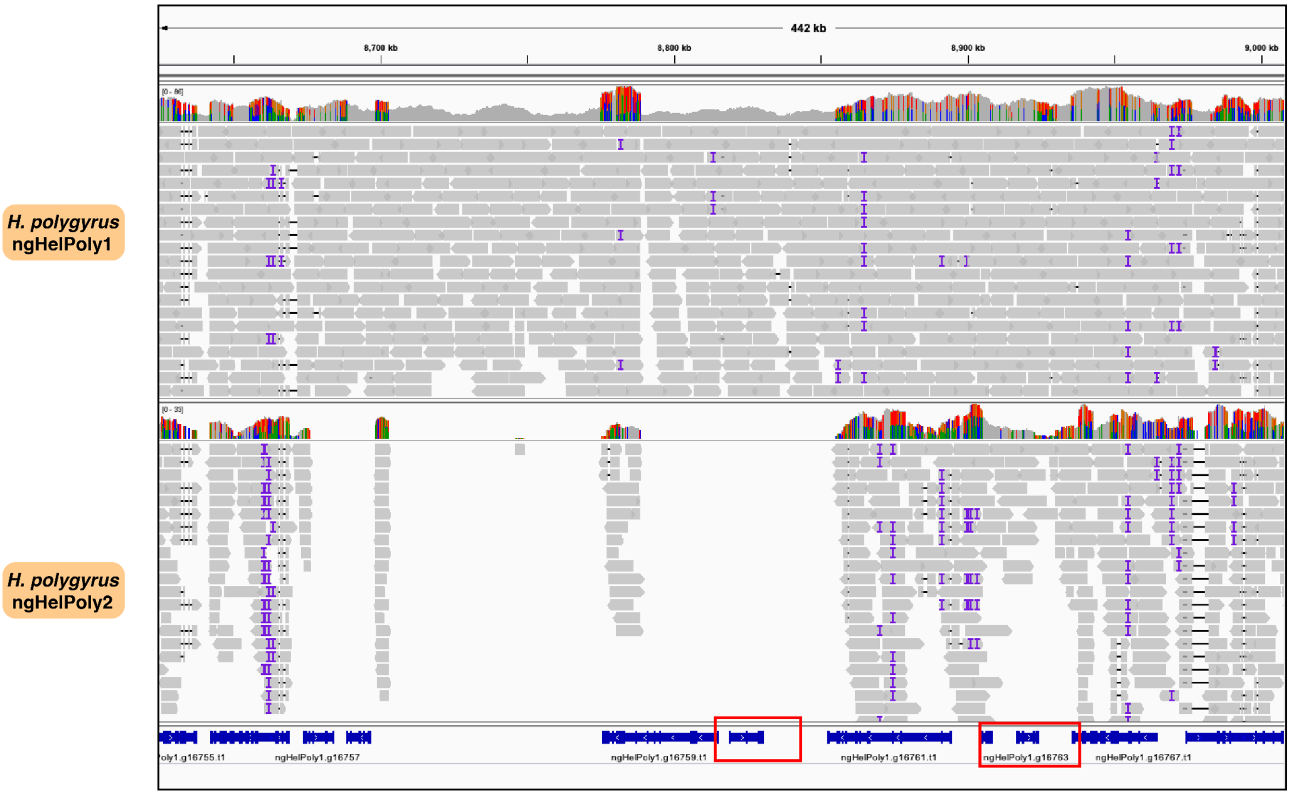


**Figure S13: Read alignments in a region in ngHelPoly1.1 containing two *Ancylostoma*-secreted proteins showing evidence of trans-specific polymorphism**
PacBio HiFi read alignments of both *H. polygyrus* individuals to the ngHelPoly1.1 reference genome in a ~383 kb region on chromosome I (I:8.62-9.01 Mb) showing evidence of trans-specific polymorphisms. Two genes in this region belong to the ASP family (ngHelPoly1.g16760.t1 and ngHelPoly1.g16763.t1; highlighted with red boxes). The top panel shows the coverage and the bottom panel shows aligned PacBio HiFi reads. Note that mismatched bases are not shown in reads at this zoom level, but can be seen as coloured vertical lines in the coverage tracks above the read alignments. This region is orthologous to the region shown in Figure 4 and Figure S11.

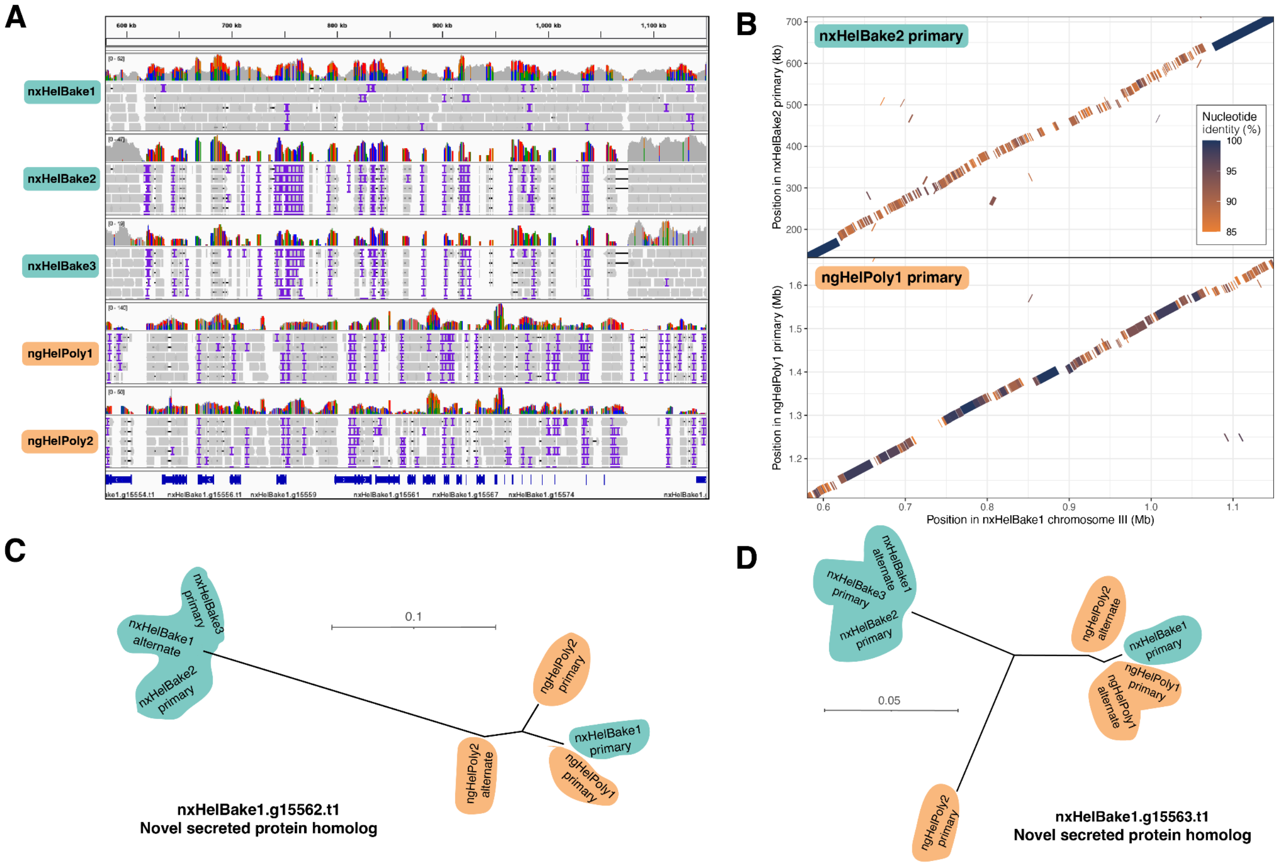


### Figure S14: Trans-specific polymorphism in novel-secreted proteins homologs

A) PacBio HiFi read alignments to the nxHelBake1.1 reference genome in a ~570 kb region on chromosome III (0.58-1.15 Mb). The top panel shows the coverage and the bottom panel shows aligned PacBio HiFi reads. The coloured vertical lines indicate mismatched bases at that position. (B) Nucleotide alignments between the alternate *H. bakeri* haplotype (represented by nxHelBake2 primary), ngHelPoly1 primary and the nxHelBake1 reference haplotype. Repetitive alignments are not shown. Gene trees of (D) nxHelBake1.g15562.t1 (E) nxHelBake1.g15563.t1 showing evidence of haplotype sharing between nxHelBake1 primary and various *H. polygyrus* haplotypes. Trees were inferred using IQ-TREE under the LG+Γ substitution model. Scale is shown in substitutions per site. Outgroup not shown.

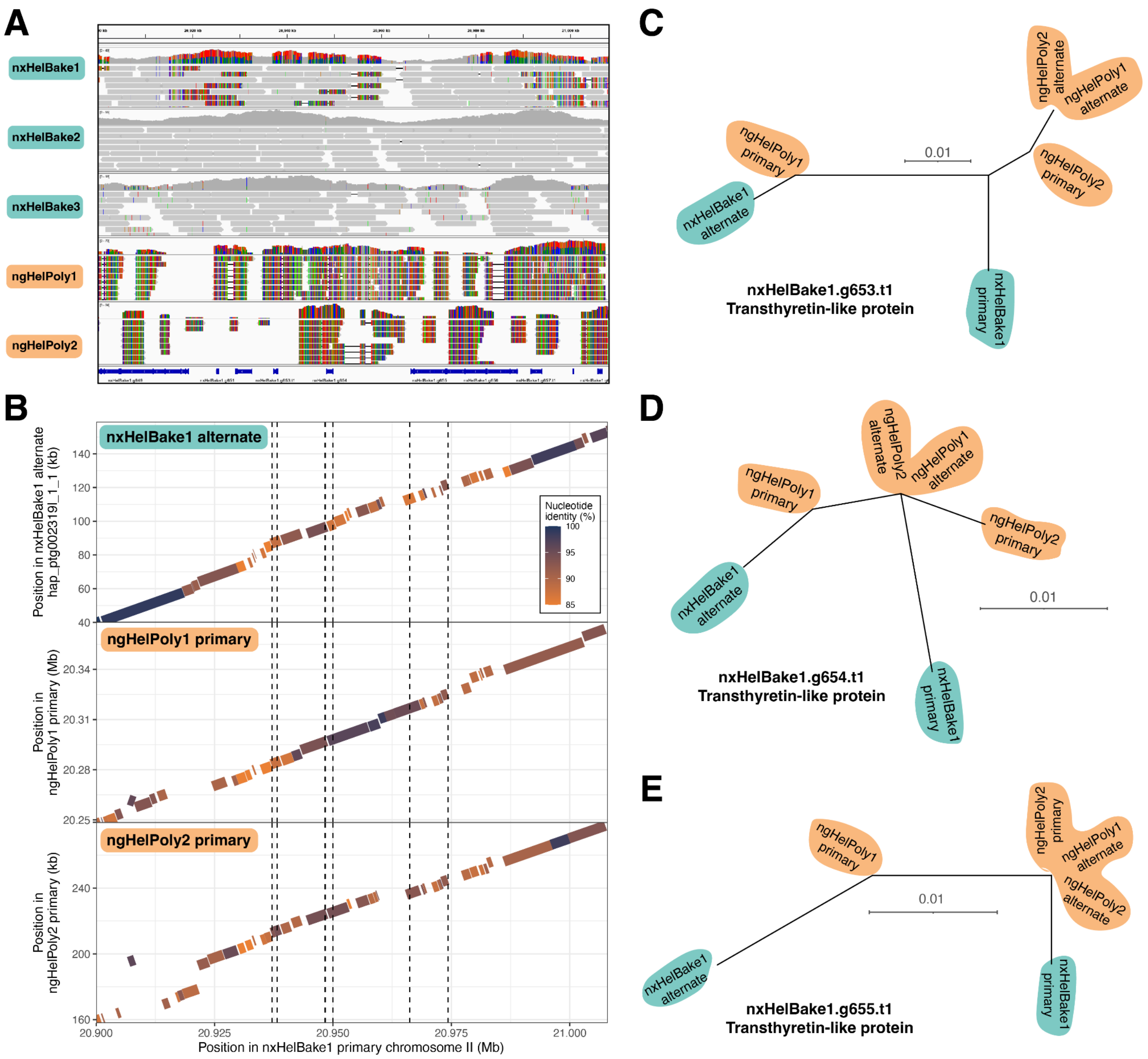


**Figure S15: Trans-specific polymorphism in a region containing multiple transthyretin-like proteins**

A) PacBio HiFi read alignments to the nxHelBake1.1 reference genome in a 177 kb region on chromosome II (20.9 - 21.1 Mb). The top panel shows the coverage and the bottom panel shows aligned PacBio HiFi reads. The coloured vertical lines indicate mismatched bases at that position. (B) Nucleotide alignments between the alternate *H. bakeri* haplotype (represented by nxHelBake1 alternate), ngHelPoly1 primary, ngHelPoly2 primary and the nxHelBake1 reference haplotype. Repetitive alignments are not shown. Gene trees of (C) nxHelBake1.g653.t1, (D) nxHelBake1.g654.t1, and (E) nxHelBake1.g655.t1 showing evidence of haplotype sharing between *H. bakeri* and *H. polygyrus* haplotypes. Tree inferred using IQ-TREE under the LG+Γ substitution model. Scale is shown in substitutions per site. Outgroup not shown.

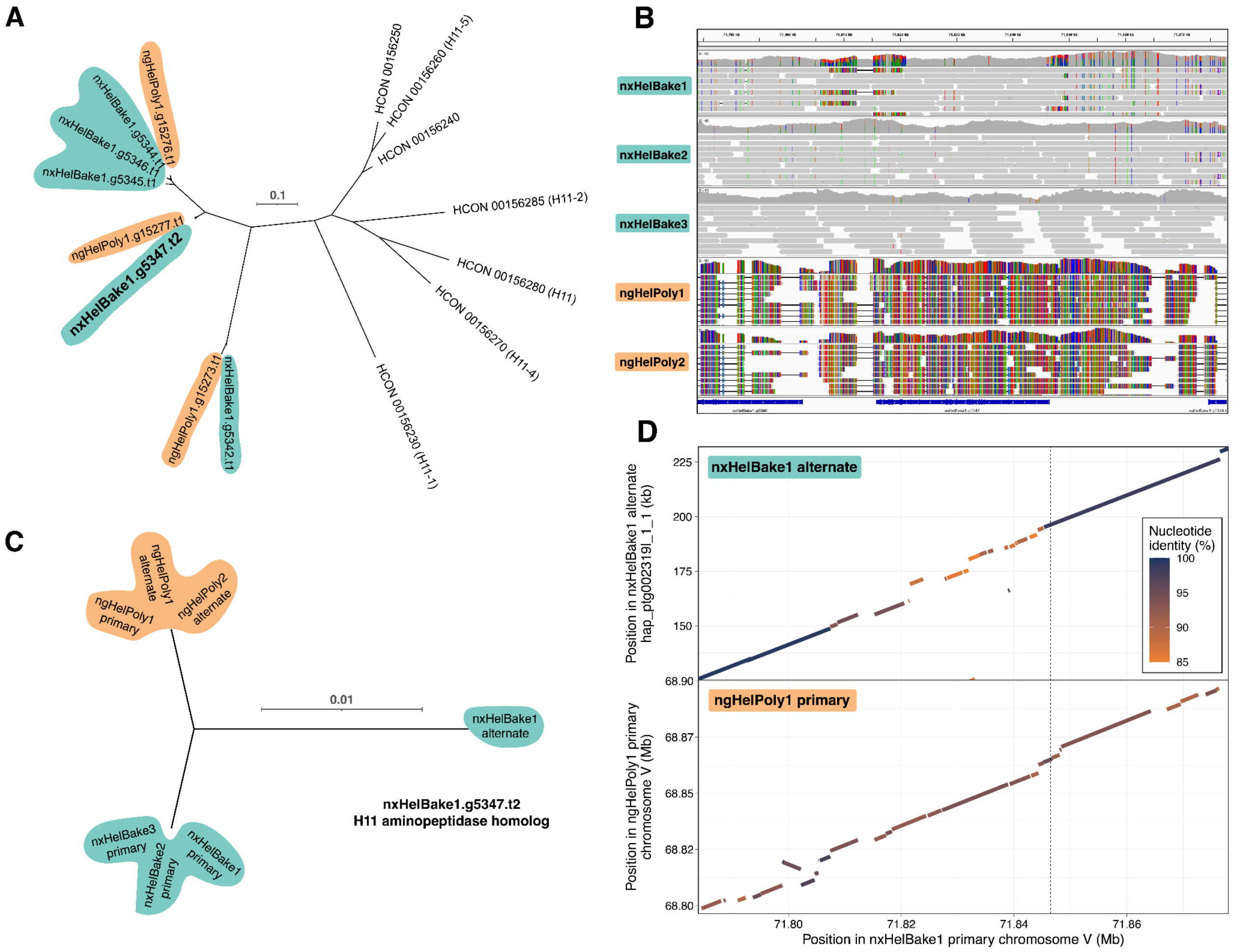


### Figure S16: Trans-specific polymorphism in a homolog of H11 aminopeptidase

A) Gene tree of H11 homologs in *H. bakeri*, *H. polygyrus* and *H. contortus*. Tree inferred using IQ-TREE under the LG+Γ substitution model. Scale is shown in substitutions per site. nxHelBake1.g5347.t2 is highlighted in bold. The names of *H. contortus* homologs are shown in parentheses which were inferred using phylogenetic relationships to previously published H11 protein sequences downloaded from NCBI. (B) PacBio HiFi read alignments to the nxHelBake1.1 reference genome in a 94 kb region on chromosome V (71.78-71.88 Mb). The top panel shows the coverage and the bottom panel shows aligned PacBio HiFi reads. The coloured vertical lines indicate mismatched bases at that position. (C) Gene tree of nxHelBake1.g5347.t2 and its homologs in the other *H. bakeri* and *H. polygyrus* haplotypes. Tree inferred using IQ-TREE under the LG+Γ substitution model. Scale is shown in substitutions per site. Outgroup not shown. (D) Nucleotide alignments between the alternate *H. bakeri* haplotype (represented by nxHelBake1 alternate), ngHelPoly1 primary and the nxHelBake1 reference haplotype. Repetitive alignments are not shown.

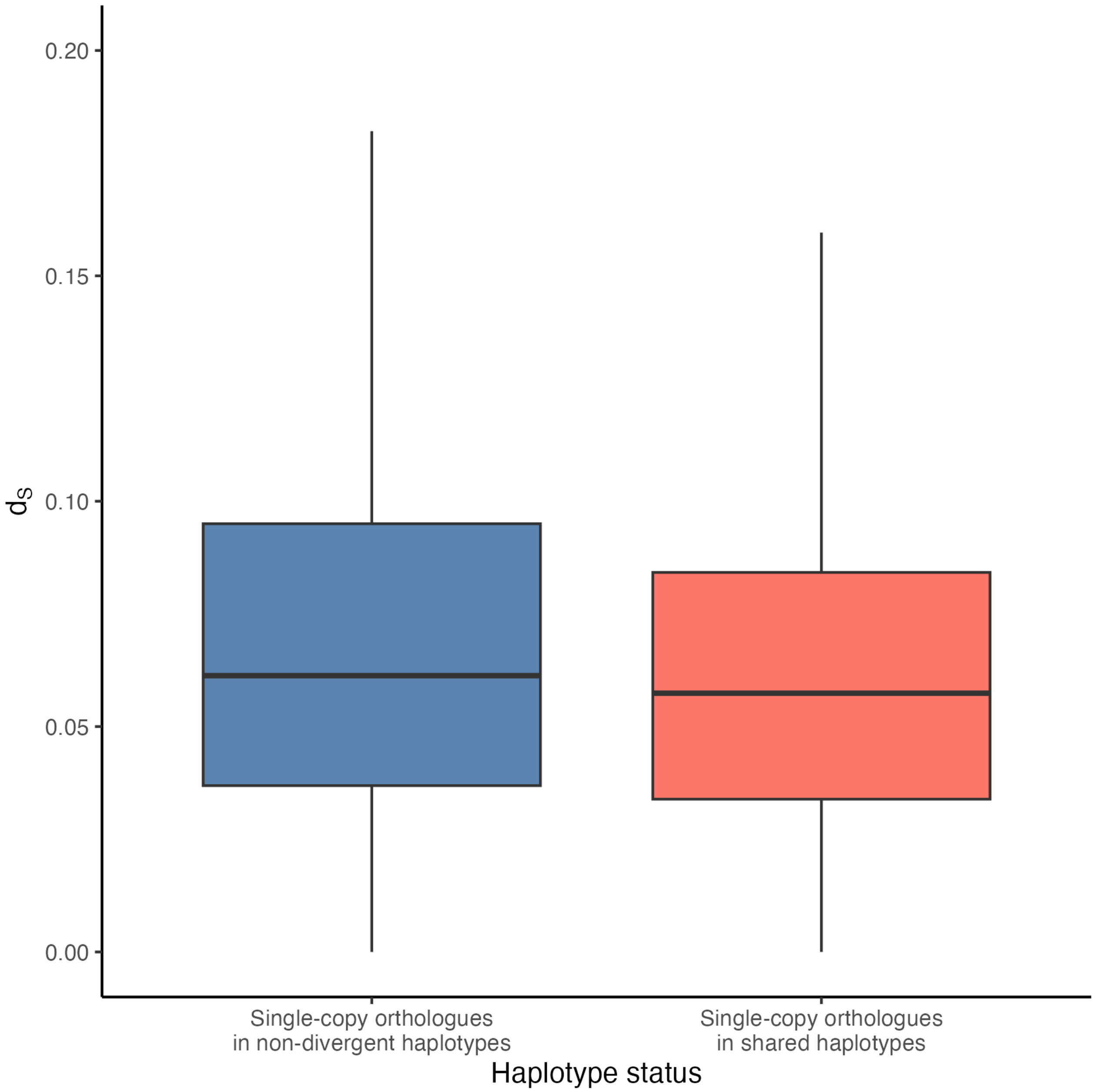


**Figure S17: Shared haplotypes show similar levels of divergence to the genome-wide average**
The average synonymous site divergence ($d_S$) for one-to-one orthologs in shared haplotypes (shared; $N$=189) and for all other orthologues (non-shared; $N$=9,753). A two-sided Wilcoxon test suggests that there is no statistical difference between the mean $d_S$ in shared and non-shared haplotypes ($p$-value = 0.1362).
